# Differential effects of human TAU isoforms: Somatic retention of 2N-TAU and increased microtubule number induced by 4R-TAU

**DOI:** 10.1101/2020.06.16.154757

**Authors:** S. Bachmann, M. Bell, J. Klimek, H. Zempel

**Affiliations:** Institute of Human Genetics and Center for Molecular Medicine Cologne (CMMC), University of Cologne, Faculty of Medicine and University Hospital Cologne

## Abstract

In the adult human brain, six isoforms of the microtubule-associated protein TAU are expressed, which result from alternative splicing of exons 2, 3 and 10 of the *MAPT* gene. These isoforms differ in the number of N-terminal inserts (0N, 1N, 2N) and C-terminal repeat domains (3R or 4R) and are differentially expressed depending on the brain region and developmental stage. Although all TAU isoforms can aggregate and form neurofibrillary tangles, some tauopathies, such as Pick’s Disease and Progressive Supranuclear Palsy, are characterized by the accumulation of specific TAU isoforms. The influence of the individual TAU isoforms in a cellular context, however, is understudied. In this report, we investigated the subcellular localization of the human-specific TAU isoforms in primary neurons, and analyzed TAU isoform-specific effects on cell area and microtubule dynamics in SH-SY5Y neuroblastoma cells. Our results show that 2N-TAU isoforms are particularly retained from axonal sorting and that axonal enrichment is independent from the number of repeat domains, but that the additional repeat domain of 4R-TAU isoform results in a general reduction of cell size and an increase of microtubule counts in cells expressing 4R-TAU isoforms. Our study points out that individual TAU isoforms may influence microtubule dynamics differentially both by different sorting patterns as well as by direct effects on microtubule dynamics.

## Introduction

The *MAPT* gene on chromosome 17 encodes the human microtubule-associated protein TAU. Expression of *MAPT* results in six major TAU isoforms in the adult human central nervous system and two isoforms in the peripheral nervous system^1–4^. The brain specific isoforms vary in the number of N-terminal inserts (0N, 1N or 2N) and C-terminal repeat domains (3R or 4R) due to alternative splicing of exons 2, 3 and 10 ^1^. TAU isoform expression is directly linked to brain development: During neurogenesis, only the shortest TAU isoform, 0N3R, is expressed, whereas in the adult brain, all six isoforms are present with roughly equal amounts of 3R- and 4R-isoforms^1,5^. Accumulation of TAU in neurofibrillary tangles (NFTs) is a hallmark of many neurodegenerative diseases, named tauopathies. All isoforms are potent to form NFTs under pathological conditions; causes can be mutations in *MAPT* affecting splicing or function of TAU, or mislocalization of TAU into the somatodendritic compartment upon cellular stress (reviewed in ^6^). Tauopathies can be classified via the isoforms that accumulate in NFTs: While TAU tangles mainly consist of 3R-TAU isoforms, e.g. in Pick’s Disease (PiD) and 4R-TAU in Progressive Supranuclear Palsy (PSP), both 3R- and 4R-TAU isoforms are present in NFTs of Alzheimer’s Disease patients^7–9^. During brain development and especially during neuronal polarization, TAU becomes efficiently sorted into the axon^10^. In the adult human brain, TAU is mainly localized in the axon, however, a small fraction can also be observed in the somatodendritic compartment and in the nucleus^11–13^. The subcellular distribution of TAU seems to be isoform-specific, e.g. 2N isoforms show a higher propensity for a somatodendritic localization than other variants^14^. Axonal targeting of TAU is thought to be mediated by a variety of processes, such as the presence of a TAU diffusion barrier (TDB) at the axon initial segment (AIS), which prevents retrograde diffusion of TAU^14,15^. Furthermore, microtubule binding affinity of TAU might be higher in the axon, likely accomplished by the presence or absence of post-translational modifications (PTMs), such as phosphorylation and acetylation^16–18^. TAU interactions are also important for proper sorting of TAU., e.g. interaction with the calcium-regulated plasma membrane– binding protein Annexin A2 was shown to link microtubules and the membrane of the growth cone, thereby trapping TAU at the presynaptic membrane^19^. Through its interaction with microtubules, TAU supports axonal differentiation, morphogenesis, outgrowth, transport and neuronal plasticity^20–22^. *In vitro* studies already described a reduced microtubule binding affinity and assembly for 3R-TAU isoforms^23–25^, which is in line with the fact that the C-terminal repeat domains together with the proline rich linker domain mediate microtubule binding^26^. If and how microtubule dynamics are altered by different TAU isoforms *in vivo* is still unclear and might depend on the differential subcellular localization of the isoforms. In this study, we address these questions by using primary mouse neurons and SH-SY5Y human neuroblastoma cells as neuronal model systems. Our results show that human TAU isoforms differ in their subcellular localization and that especially 2N-TAU isoforms are less axonally enriched than shorter isoforms. In addition, we show an isoform-specific effect on cell size and microtubule number: Expression of 4R-TAU isoforms, for example, resulted in smaller cells and increased microtubule counts. Microtubule dynamics, such as microtubule stability, length, and growth rate, were not altered upon expression of different TAU isoforms in undifferentiated SH-SY5Y cells. We show here that individual TAU isoforms may influence microtubule dynamics differentially both by different sorting patterns as well as by direct effects on microtubule dynamics in a non-disease background.

## Methods

### Cell culture

SH-SY5Y cells were cultured in DMEM/F12, GlutaMAX (Thermofisher Scientific (TFS)) supplemented with 10% Fetal Bovine Serum (FBS, Biochrom AG) and 1× Antibiotic-/Antimycotic solution (TFS) in a humidified incubator at 37°C, 5% CO^2^. Primary mouse neurons were isolated and cultured as described before^27^ with slight modifications. In brief, brains of FVB/N mouse embryos were dissected at embryonic day 13.5. Brainstem and meninges were removed and whole cortex was digested with 1× Trypsin (Panbiotech). Neurons were diluted in pre-warmed (37°C) neuronal plating medium (Neurobasal media (TFS), 1% FBS, 1× Antibiotic-/Antimycotic solution (TFS), 1× NS21 (Panbiotech)) and cultivated in a humidified incubator at 37°C, 5% CO_2_. Four days after plating, media was doubled with neuronal maintenance media (Neurobasal media (TFS)), 1× Antibiotic-/Antimycotic solution (TFS), 1× NS21 (Panbiotech)) and cells were treated with 0.5 µg/ml AraC (Sigma-Aldrich).

### Microtubule dynamics

SH-SY5Y cells were co-transfected with expression plasmids containing Dendra2c-TAU (TAU^D2^) isoforms and tdTomato-N1-EB3 by using Lipofectamine2000 (TFS) according the manufacturers protocol. Control cells were transfected with tdTomato-N1-EB3 and an empty Dendra2c plasmid. Only cells showing both, Dendra2c and tdTomato signal, were used for analysis. Two days after transfection, cells were transferred into a live-cell imaging chamber (ALA Scientific) and EB3 comets of a single cell were imaged for 120 seconds (1 frame per second) with a Leica DMi8 microscope (Leica). Original movie file was processed by ImageJ software (Schindelin *et al*., 2012; Schneider, Rasband and Eliceiri, 2012) as follows: An average minimum projection was calculated from all frames and subsequently subtracted from all. Afterwards threshold was set and microtubule dynamics were analyzed by ImageJ plugin TrackMate^28^, using LoG detector with an estimated blob size of 2 px. The following parameters were examined: Cell area [µm^2^], Microtubule number/µm^2^, microtubule length [µm], microtubule stability [s] and microtubule growth rate [µm/s]. Data was filtered for quality ≥1.5, track start >0 seconds and track end <120 seconds. All experiments were conducted in three replicates and at least five cells were analyzed per condition. Cell area was measured from at least 30 cells. To test for statistical significance, one-way ANOVA with correction for multiple comparisons (Tukey’s test) was performed using GraphPad Prism software.

### Cell lysis and Western blot analysis

SH-SY5Y cells were lysed with RIPA buffer (50 mM HEPES pH 7.6, 150 mM NaCl, 1 mM EDTA, 1% Triton X-100, 0.1% Sodium dodecylsulfate, 0.5% Sodium deoxycholate, 1× cOmplete ULTRA EDTA-free protease inhibitor cocktail (Merck)). Samples were diluted with SDS-buffer and separated on 10% polyacrylamide gels. Proteins were transferred to PVDF membranes and blocked in 5% non-fat dry milk in TBS-T. Membranes were incubated with first antibody overnight at 4°C, washed with TBS-T and incubated with secondary HRP-coupled antibody for one hour at room temperature. Used antibodies are listed in table 1. Luminescent signals were detected with ChemiDoc XRS+ system (Bio-Rad).

**Table 1.** Antibodies used in this study.

| Antibody | Company | Cat.No. | Application | Dilution |
| --- | --- | --- | --- | --- |
| **Primary antibodies** |  |  |  |  |
| *Rabbit polyclonal anti-TAU (K9JA)* | Dako | A0024 | WB<br>ICC | 1:10000<br>1:1000 |
| *Chicken polyclonal anti-MAP2* | Abcam | ab5392 | ICC | 1:2000 |
| *Mouse monoclonal anti-HA (16B12)* | Biolegend | 901501 | ICC | 1:1000 |
| *Mouse monoclonal anti-GAPDH (1D4)* | Novus<br>Biologicals | NB300-221 | WB | 1:2000 |
| **Secondary antibodies** |  |  |  |  |
| *Donkey anti-mouse IgG AlexaFluor-488* | Thermofisher<br>Scientific | A21202 | ICC | 1:1000 |
| *Goat anti-chicken IgG AlexaFluor-647* | Thermofisher<br>Scientific | A21449 | ICC | 1:1000 |
| *Donkey anti-rabbit IgG AlexaFluor-488* | Thermofisher Scientific | A21206 | ICC | 1:1000 |
| *Phalloidin, AlexaFluo-568* | Thermofisher Scientific | A12380 | ICC | 1:40 |
| *Goat Anti-Mouse IgG (H+L), HRP-linked* | Dianova | 115-035-003 | WB | 1:20.000 |
| *Goat anti-rabbit IgG, HRP-linked* | Cell Signaling Technologies | 7074 | WB | 1:20.0000 |

### Axonal enrichment factor

To analyze TAU axonal enrichment, mouse primary neurons were transfected as described before with Lipofectamine2000 (TFS)^31^. In brief, neurons were co-transfected with the corresponding HA-tagged TAU (TAU^HA^) isoform and tdTomato. After two days, neurons were fixed and stained for endogenous mouse Tau with a panTAU antibody, or HA for exogenous TAU^HA^ and MAP2 as described before^27^, using antibodies listed in table 1. After imaging on a fluorescence microscope (Axioscope 5, Zeiss), axonal sorting was analyzed by measuring mean fluorescence intensities (MFI) of tdTomato and TAU in the axon and the soma. Axons were identified by absence of MAP2 and presence of TAU signal, branching pattern of ∼90°, a length of above 300 μm and constant diameter. Axonal enrichment factor (AEF) was calculated as follows: AEF = (MFI_(TAU,Axon)_/ MFI_(TAU,Soma)_)/ (MFI_(tdTom,Axon)_/ MFI_(tdTom,Soma)_). The experiment was performed in three replicates and at least ten cells were analyzed per condition. To test for statistical significance, one-way ANOVA with correction for multiple comparisons (Tukey’s test) was performed using GraphPad Prism software.

### Antibody list

## Results

### TAU isoforms differ in their subcellular localization and influence axonal volume

TAU is considered an axonal protein in mature neurons^11^. However, previous results indicate a difference in axonal enrichment of the six human TAU isoforms, that were overexpressed as Dendra2c-tagged fusion proteins in primary neurons^14^. To further investigate TAU isoform axonal sorting and rule out the influence of a big protein tag (such as Dendra2c with approx. 26 kDa) on cellular localization, all six human TAU isoforms were fused to an HA-tag (TAU^HA^) and co-transfected for 2 days with a volume marker (tdTomato) into primary mouse neurons aged for 7-9 days. Control neurons were transfected only with tdTomato as a volume marker, for which we assume an unbiased axodendritic distribution after two days of expression. To quantify axonal targeting of the different versions of TAU, we normalized the axonal presence of TAU against the unbiased distribution of tdTomato, expressed as the axonal enrichment factor (AEF, see methods for details). To distinguish axons and dendrites, neurons were fixed and stained for endogenous mouse TAU or transfected human TAU^HA^ using an anti-HA antibody, and the dendritic marker MAP2. Axons were identified by absence of MAP2, presence of TAU or HA signal and further morphological criteria (see methods for details; Fig. 1A). Differences in axonal sorting of TAU were observed for the different TAU^HA^ isoforms (Fig. 1B,C). For endogenous TAU, a strong axonal localization is visible from the immunofluorescence images, indicated by a strong Tau signal in the axon and only low levels in the soma and dendrites (AEF of ∼10, Fig. 1B,C). Strong axonal enrichment was also observed in neurons expressing 0N3R-TAU^HA^. In contrast, neurons expressing 2N4R-TAU^HA^ show lower levels of axonal TAU, but are still targeted to the axon compared to the presumably unbiased tdTomato distribution (Fig. 1B). To quantitatively compare the different TAU^HA^ isoforms, axonal enrichment of TAU was calculated from soma-to-axon ratio of TAU fluorescence intensity and normalized to soma-to-axon ratio of the tdTomato signal, which is assumed to be random (Fig. 1C). Axonal enrichment of endogenous Tau was approximately 10-fold higher compared to tdTomato, while the human TAU^HA^ isoforms showed approximately 4-to 6-fold enrichment (Fig. 1C). From all six transfected isoforms, 0N3R TAU^HA^ showed the strongest axonal enrichment, which was still approximately 35% less than that of endogenous TAU. A relatively efficient (but still ∼40% less than endogenous Tau) axonal sorting could also be observed for 1N3R-TAU^HA^. Of note, both 2N-TAU^HA^ isoforms showed the weakest axonal enrichment. These results indicate that efficiency of axonal sorting of TAU is isoform dependent, and might be also dependent on the number of N-terminal inserts, since 2N4R- and 2N3R-TAU^HA^ showed the lowest enrichment factors. No significant difference was observed between the other 3R- and 4R-expressing neurons, hinting towards a repeat-independent sorting mechanism of the isoforms.

**Figure 1:**
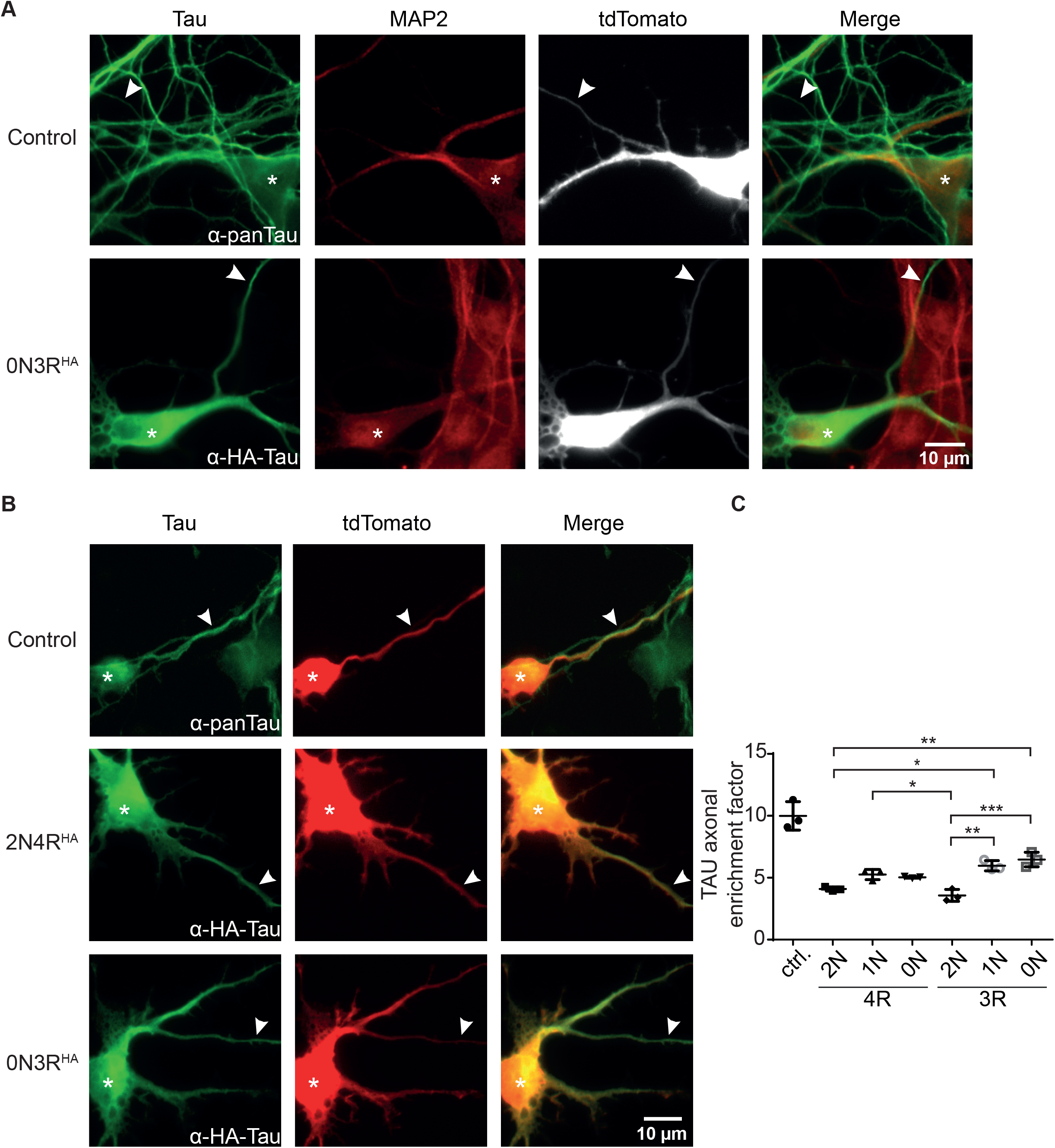
2N-containing TAU^HA^ isoforms are less efficiently sorted into the axon in mouse primary neurons. HA-tagged TAU isoforms (0N, 1N, 2N and 3R or 4R) and tdTomato as a volume marker were co-transfected into primary mouse neurons (DIV4) and expressed for two days. **(A)** Representative pictures of mouse primary neurons (DIV6), transfected with tdTomato (ctrl.) or co-transfected with tdTomato and 0N3R-TAU, respectively. Neurons were stained with α-panTAU (for endogenous TAU in control) or α-HA antibody (for transfected TAU^HA^) and α-MAP2 to distinguish axons and dendrites. Arrowheads indicate axons, asterisks indicate cell body. **(B)** Representative pictures of mouse primary neurons (DIV6), transfected with tdTomato (Ctrl.) or co-transfected with tdTomato and 2N4R-TAU^HA^ and 0N3R-TAU^HA^, respectively. Neurons were stained with α-Tau (for control) or α-HA antibody to analyze axonal enrichment of transfected or endogenous TAU. Arrowheads indicate axons, asterisks indicate cell body. **(C)** Axonal enrichment of TAU was calculated from soma-to-axon ratio of TAU fluorescence intensity and normalized to soma-to-axon ratio of the tdTomato signal. An axonal enrichment of 1 is considered as a random distribution. N=3, at least 10 cells were analyzed per condition. Error bars represent SEM. Statistical analysis was performed by one-way ANOVA with Tukey’s test for correction of multiple comparisons. Statistical significance: * = p ≤ 0.05; ** = p ≤ 0.01; *** = p ≤ 0.001.

### Differential effect of TAU isoforms on cell size and microtubule number

One main function of TAU is its role in microtubule stability and spacing^32,33^. The influence of individual TAU isoforms on microtubule dynamics, however, has not been addressed in living cells. To investigate TAU isoform specific effects on microtubules, SH-SY5Y neuroblastoma cells were transfected with TAU^D2^ isoforms. Expression levels of TAU^D2^ were confirmed by Western blot analysis (Fig. 2A). Of note, only low levels of endogenous TAU were observed in control and transfected cells (Fig. 2A long exposure). Since microtubules are essential components of the cytoskeleton, initial analysis of TAU isoform-specific effects focused on changes in cell size, which may hint to cytoskeletal alterations. Cell area was significantly (∼20%) smaller in cells overexpressing 4R-TAU^D2^ isoforms compared control cells (Fig. 2B). In addition, cells expressing 2N3R- and 0N3R-TAU^D2^ had a significantly greater cell area then 4R expressing cells (∼20%), comparable to untransfected cells, while 1N3R only showed a trend towards increased cell size compared to the 4R-TAU^D2^-isoforms expressing cells (∼15% bigger cell size). To further investigate the role of TAU isoforms in microtubule dynamics, SH-SY5Y cells were co-transfected with the corresponding TAU^D2^ isoform and tdTomato-N1-EB3 (Fig. 2C-G). EB3 binds the growing microtubule plus-ends and can be used in live-cell imaging to track microtubule assembly^34^. Expression of TAU^D2^ was confirmed via the green fluorescence of the D2-tag in life imaging. EB3 comets were recorded for two minutes (1 frame per second), images were processed as described in Fig. 2C and microtubule dynamics were analyzed afterwards using TrackMate plug-in for ImageJ^28^. Overall number of microtubules were quantified and normalized to the corresponding cell area (Fig. 2D). Microtubule count was significantly increased compared to control cells (5.5 MTs/ µm^2^) in cells expressing 2N4R- and 0N4R-TAU^D2^ (7.2 MTs/ µm^2^ and 7.3 MTs/ µm^2^ respectively). In contrast, 3R-TAU^D2^ isoforms showed no difference in microtubule counts compared to control cells, despite a slight trend towards a reduced number of microtubules for 0N3R. Comparing 3R- and 4R-TAU^D2^ isoforms showed that microtubule numbers were significantly decreased for 0N3R compared to 4R-TAU^D2^ expressing cells (Fig. 2D). To further investigate, how microtubule counts might be affected by TAU^D2^ isoforms, microtubule stability over the recorded time, total microtubule length and microtubule growth rate (Fig. 2E-G) were analyzed. We observed several tendencies that did, however, not reach statistical significance: 1N- and 0N-TAU^D2^ isoforms seem to cause a slight decrease in microtubule stability independent of the presence of the second N-terminal repeat (Fig. 2E), and 2N4R-TAU^D2^ seems to increase microtubule length (Fig 2F). Microtubule growth rate is slightly, but not significantly, increased for individual TAU isoform overexpressing cells compared to control cells (Fig. 2G), but significantly increased (p<0.0143, unpaired t-test), when we compared all TAU isoform expressing cells to control cells. All in all, the results indicate that TAU isoforms have a significant impact on cell size and on microtubule counts, without significant differential effects on microtubule stability, length, and growth rate in undifferentiated SH-SY5Y cells.

**Figure 2:**
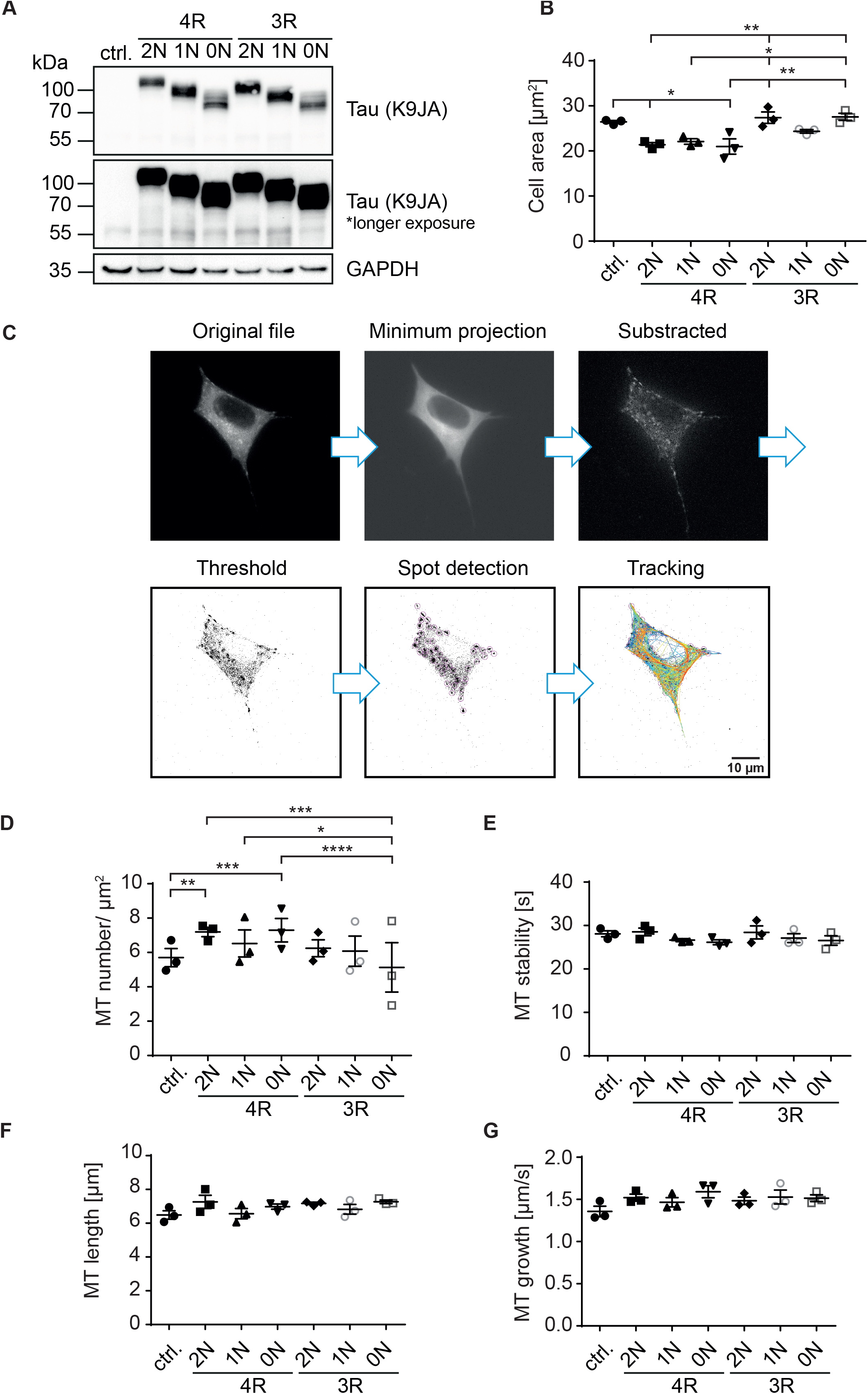
4R-TAU isoforms decrease cell size and increase microtubule counts in undifferentiated SH-SY5Y neuroblastoma cells. Expression of TAU^D2^ isoforms (0N, 1N, 2N and 3R or 4R) in undifferentiated SH-SY5Y cells.**(A)** Western blot of SH-SY5Y cells transfected with the corresponding TAU^D2^ isoforms. Longer exposure of TAU signal shows negligible endogenous TAU expression (∼55 kDa). GAPDH was used as a loading control. **(B)** Cell area was measured from SH-SY5Y cells, co-transfected with tdTomato-N1-EB3 and TAU^D2^. Cell area was analyzed from at least 30 cells. Error bars represent SEM. Statistical analysis was performed by one-way ANOVA with Tukey’s test for correction of multiple comparisons. Asterisks indicate statistical significance: * = p ≤ 0.05; ** = p ≤ 0.01. **(C)** Image processing for input into TrackMate. Minimum projection was calculated from all frames and subtracted from all. Afterwards threshold was set, and microtubule dynamics were analyzed by TrackMate. **(D-F)** Microtubule dynamics of SH-SY5Y cells co-transfected with Dendra2c-tagged TAU isoforms (0N, 1N, 2N and 3R or 4R) and tdTomato-N1-EB3. Growing microtubule plus-ends were monitored in living cells for two minutes (1 frame per second). The following parameters were examined: **(D)** Microtubule number [MT number/µm2] (normalized to corresponding cell area), **(E)** microtubule length [µm], **(F)** microtubule stability [s] and **(G)** microtubule growth rate [µm/s.] N=3, at least 5 cells were analyzed per condition. Error bars represent SEM. Statistical analysis was performed by one-way ANOVA with Tukey’s test for correction of multiple comparisons. Asterisks indicate statistical significance: * = p ≤ 0.05; ** = p ≤ 0.01; *** = p ≤ 0.001; **** = p ≤ 0.0001.

## Discussion

In this study, we show that human TAU isoforms can influence microtubule dynamics in a differential matter, though i) differential compartment-specific cellular distribution, and ii) differential effects on cell size and microtubule count/density, but to a lesser extent on microtubule dynamics. Expressed TAU-isoforms with an N-terminal HA-tag differ in their subcellular localization in primary mouse neurons already after two days of expression. Especially 2N-TAU isoforms showed less axonal enrichment than shorter isoforms, which is in line with previous findings, demonstrating less axonal enrichment for Dendra2c-tagged 2N-TAU isoforms^14^. The shortest human TAU isoform, 0N3R, showed the strongest axonal enrichment compared to the other isoforms. Especially 2N-TAU isoforms poorly localize to the axon. This might be due to their increased size compared to other TAU isoforms, although the 2N3R and 1N4R have nearly the same size, with 410 and 412 aa respectively. No significant difference was observed between the other 3R and 4R isoform expressing neurons, pointing towards a repeat-independent sorting mechanism of the isoforms. Axonal sorting of TAU is also influenced by PTMs, such as phosphorylation and acetylation, which might be different for the TAU isoforms, however this has not been investigated in detail for isoform-specific effects^16–18^. Of note, endogenous mouse TAU axonal sorting is more efficient compared to all HA-tagged human TAU isoforms. This might have methodological reasons: 1.) Expression time was relatively short in our study, increasing it from two days to four days might be beneficial for the axonal enrichment of exogenous TAU. 2.) Expression levels of the individual isoforms compared to endogenous TAU were found to be roughly ∼20 fold higher in previous studies using similar techniques^15^, it is thus possible that the sorting machinery is simply saturated due to the high (and relatively short) expression. 3.) The HA-tag might impair efficient axonal targeting of endogenous TAU, as seen also for Dendra2c-tagged Tau^14^, due to its molecular size or by interfering with critical TAU interactions. 4.) Sorting of human TAU might be different from mouse TAU, which could have several reasons: Young primary neurons, aged roughly one week, mainly express 0N3R-TAU, mouse TAU in general is slightly shorter, and protein lifetime might be different due to tagging or changed protein interactions^1,35^. Overall, we show here that in polarized primary neurons, human TAU isoforms differ remarkably in their axonal sorting efficiencies (with 0N3R-TAU sorting 1.5x better than the 2N-TAU isoforms), and that the somatic retention of 2N-TAU isoforms is repeat domain independent.

To study isoform-specific effects of TAU on microtubule dynamics in a cellular context, we used the well-established SH-SY5Y neuroblastoma cell line, which expresses only low levels of endogenous Tau in the undifferentiated state^36^. 4R-TAU isoforms expressing cells showed a significant reduction in cell size (∼30%); however, microtubule numbers were increased in these cells. These results go in line with *in vitro* studies showing an increased binding affinity and assembly of microtubules by 4R-TAU isoforms^23–25^. Since cell size is affected in undifferentiated SH-SY5Y cells, there might be also an isoform-specific effect on neurite growth, which needs to be investigated in differentiated neuronal models. Microtubule dynamics were not significantly altered by the expression of individual TAU isoforms in our model system, however there might be a compartment-specific effect visible in neurons, due to the differential localization of TAU isoforms demonstrated in the first part of this study. Possible compartment-specific effects might be caused by different PTMs, differences in microtubule binding affinity and different protein-protein interactions of TAU isoforms. Of note, our results suggest that the N-terminal inserts play a major role in subcellular localization of TAU, and thus may be important for the development or prevention of pathological processes, such as missorting of TAU into the somatodendritic compartment and subsequent synapse loss. This is in line with the observation, that presence of the H2 haplotype of the *MAPT* locus, resulting in a two times lower expression of exon 3+ transcripts (2N-TAU isoforms respectively), has neuroprotective effects^37,38^. 2N-TAU isoforms were also shown to differ in their interactome from the other TAU isoforms, suggesting differences in the cellular functions of the TAU isoforms^39^. While the generation and effects of the 3R- and 4R-TAU isoforms have been studied in detail, splicing and behavior of the N-terminally different isoforms are understudied. Future studies with polarized cells, such as primary and iPSC-derived neurons, should aim to further characterize TAU isoforms regarding their impact on microtubule dynamics in different subcellular compartments, taking into account the differential sorting, impact on cell size and microtubule number. In sum, we showthat i) the efficiency of human TAU sorting into the axon is isoform-dependent, with 2N-isoforms being most retained in the somatodendritic compartment, and ii) 4R-TAU isoforms result in a general reduction of cell size and an increase of microtubule counts. This points to isoform- and compartment-specific functions of TAU in neurons. Follow-up experiments are needed to clarify how the different TAU isoforms might influence cellular functions and if they contribute differentially to the pathological processes underlying AD and related tauopathies.

## Acknowledgements

We thank Prof. R. Wiesner (Institute for Vegetative Physiology, University Hospital Cologne) for providing SH-SY5Y cells. Live-cell imaging experiments were performed at CMMC microscopy facility (Cologne, Germany). Animals were provided by CMMC animal facility (Cologne, Germany) and CECAD in vivo research facility (Cologne, Germany).

## Author contributions

SB: study design, data acquisition, analysis, and interpretation, drafting of manuscript. MB and JK: assistance in data acquisition and methodology development. MB: manuscript proofreading. HZ: project funding, providing of concept, study design, interpretation of data, drafting of manuscript.

## References

1. Goedert, M., Spillantini, M. G., Jakes, R., Rutherford, D. & Crowther, R. A. Multiple isoforms of human microtubule-associated protein tau: sequences and localization in neurofibrillary tangles of Alzheimer’s disease. Neuron (1989) doi:10.1016/0896-6273(89)90210-9.

2. Andreadis, A., Brown, W. M. & Kosik, K. S. Structure and novel exons of the human .tau. gene. Biochemistry 31, 10626–10633 (1992).

3. Couchie, D. et al.. Primary structure of high molecular weight tau present in the peripheral nervous system. Proc. Natl. Acad. Sci. 89, 4378–4381 (1992).

4. Goedert, M., Spillantini, M. G. & Crowther, R. A. Cloning of a big tau microtubule-associated protein characteristic of the peripheral nervous system. Proc. Natl. Acad. Sci. 89, 1983–1987 (1992).

5. Trabzuni, D. et al.. MAPT expression and splicing is differentially regulated by brain region: relation to genotype and implication for tauopathies. Hum. Mol. Genet. 21, 4094–4103 (2012).

6. Arendt, T., Stieler, J. T. & Holzer, M. Tau and tauopathies. Brain Res. Bull. 126, 238–292 (2016).

7. Goedert, M. et al.. Molecular dissection of the paired helical filament. Neurobiol. Aging 16, 325–334 (1995).

8. Buee, L. & Delacourte, A. Comparative biochemistry of tau in progressive supranuclear palsy, corticobasal degeneration, FTDP-17 and Pick’s disease. Brain Pathol. 9, 681–693 (1999).

9. Arai, T. et al. Different immunoreactivities of the microtubule-binding region of tau and its molecular basis in brains from patients with Alzheimer’s disease, Pick’s disease, progressive supranuclear palsy and corticobasal degeneration. Acta Neuropathol. 105, 489–498 (2003).

10. Mandell, J. W. & Banker, G. A. The microtubule cytoskeleton and the development of neuronal polarity. Neurobiol. Aging (1995) doi:10.1016/0197-4580(94)00164-V.

11. Binder, L. I., Frankfurter, A. & Rebhun, L. I. The distribution of tau in the mammalian central nervous system. J. Cell Biol. 101, 1371–1378 (1985).

12. Rady, R. M., Zinkowski, R. P. & Binder, L. I. Presence of tau in isolated nuclei from human brain. Neurobiol. Aging 16, 479–486 (1995).

13. Tashiro, K., Hasegawa, M., Ihara, Y. & Iwatsubo, T. Somatodendritic localization of phosphorylated tau in neonatal and adult rat cerebral cortex. Neuroreport 8, 2797–2801 (1997).

14. Zempel, H. et al. Axodendritic sorting and pathological missorting of Tau are isoform-specific and determined by axon initial segment architecture. J. Biol. Chem. 292, 12192–12207 (2017).

15. Li, X. et al. Novel diffusion barrier for axonal retention of Tau in neurons and its failure in neurodegeneration. EMBO J. 30, 4825–4837 (2011).

16. Evans, D. B. et al. Tau Phosphorylation at Serine 396 and Serine 404 by Human Recombinant Tau Protein Kinase II Inhibits Tau’s Ability to Promote Microtubule Assembly. J. Biol. Chem. 275, 24977–24983 (2000).

17. Kishi, M., Pan, Y. A., Crump, J. G. & Sanes, J. R. Mammalian SAD kinases are required for neuronal polarization. Science (80-.). (2005) doi:10.1126/science.1107403.

18. Tsushima, H. et al. HDAC6 and RhoA are novel players in Abeta-driven disruption of neuronal polarity. Nat Commun 6, 7781 (2015).

19. Gauthier-Kemper, A. et al. The frontotemporal dementia mutation R406W blocks tau’s interaction with the membrane in an annexin A2-dependent manner. J. Cell Biol. (2011) doi:10.1083/jcb.201007161.

20. Esmaeli-Azad, B., McCarty, J. H. & Feinstein, S. C. Sense and antisense transfection analysis of tau function: Tau influences net microtubule assembly, neurite outgrowth and neuritic stability. J. Cell Sci. (1994).

21. Kempf, M., Clement, A., Faissner, A., Lee, G. & Brandt, R. Tau binds to the distal axon early in development of polarity in a microtubule- and microfilament-dependent manner. J. Neurosci. (1996).

22. Takei, Y., Teng, J., Harada, A. & Hirokawa, N. Defects in axonal elongation and neuronal migration in mice with disrupted tau and map1b genes. J. Cell Biol. 150, 989–1000 (2000).

23. Goedert, M. & Jakes, R. Expression of separate isoforms of human tau protein: correlation with the tau pattern in brain and effects on tubulin polymerization. EMBO J. 9, 4225–4230 (1990).

24. Goode, B. L., Chau, M., Denis, P. E. & Feinstein, S. C. Structural and functional differences between 3-repeat and 4-repeat tau isoforms. Implications for normal tau function and the onset of neurodegenetative disease. J. Biol. Chem. 275, 38182–38189 (2000).

25. Panda, D., Samuel, J. C., Massie, M., Feinstein, S. C. & Wilson, L. Differential regulation of microtubule dynamics by three- and four-repeat tau: Implications for the onset of neurodegenerative disease. Proc. Natl. Acad. Sci. 100, 9548 LP – 9553 (2003).

26. Goode, B. L. et al. Functional interactions between the proline-rich and repeat regions of tau enhance microtubule binding and assembly. Mol. Biol. Cell 8, 353– 365 (1997).

27. Zempel, H. & Mandelkow, E. M. Tracking Tau in neurons: How to grow, fix, and stain primary neurons for the investigation of Tau in all developmental stages. in Methods in Molecular Biology (2017). doi:10.1007/978-1-4939-6598-4_20.

28. Tinevez, J.-Y. et al. TrackMate: An open and extensible platform for single-particle tracking. Methods 115, 80–90 (2017).

29. Schindelin, J. et al. Fiji: an open-source platform for biological-image analysis. Nat. Methods 9, 676–682 (2012).

30. Schneider, C. A., Rasband, W. S. & Eliceiri, K. W. NIH Image to ImageJ: 25 years of image analysis. Nat Methods 9, 671–675 (2012).

31. Zempel, H., Luedtke, J. & Mandelkow, E. M. Tracking Tau in neurons: How to transfect and track exogenous tau into primary neurons. in Methods in Molecular Biology (2017). doi:10.1007/978-1-4939-6598-4_21.

32. Weingarten, M. D., Lockwood, A. H., Hwo, S. Y. & Kirschner, M. W. A protein factor essential for microtubule assembly. Proc. Natl. Acad. Sci. U. S. A. (1975) doi:10.1073/pnas.72.5.1858.

33. Witman, G. B., Cleveland, D. W., Weingarten, M. D. & Kirschner, M. W. Tubulin requires tau for growth onto microtubule initiating sites. Proc. Natl. Acad. Sci. U. S. A. 73, 4070–4074 (1976).

34. Stepanova, T. et al. Visualization of Microtubule Growth in Cultured Neurons via the Use of EB3-GFP (End-Binding Protein 3-Green Fluorescent Protein). J. Neurosci. 23, 2655 LP – 2664 (2003).

35. Kosik, K. S., Orecchio, L. D., Bakalis, S. & Neve, R. L. Developmentally regulated expression of specific tau sequences. Neuron 2, 1389–1397 (1989).

36. Bell, M. & Zempel, H. SH-SY5Y-derived Neurons: A Neuronal Model System for Investigating TAU Sorting Mechanisms and Neuronal Subtype-specific TAU Vulnerability. Preprints (2020) doi:10.20944/preprints202006.0203.v1.

37. Beevers, J. E. et al. MAPT Genetic Variation and Neuronal Maturity Alter Isoform Expression Affecting Axonal Transport in iPSC-Derived Dopamine Neurons. Stem Cell Reports 9, 587–599 (2017).

38. Caffrey, T. M., Joachim, C., Paracchini, S., Esiri, M. M. & Wade-Martins, R. Haplotype-specific expression of exon 10 at the human MAPT locus. Hum. Mol. Genet. 15, 3529–3537 (2006).

39. Liu, C., Song, X., Nisbet, R. & Götz, J. Co-immunoprecipitation with Tau Isoform-specific Antibodies Reveals Distinct Protein Interactions and Highlights a Putative Role for 2N Tau in Disease. J. Biol. Chem. 291, 8173–8188 (2016).

